# A miniaturized threshold-triggered acceleration data-logger for recording burst movements of aquatic animals

**DOI:** 10.1101/203828

**Authors:** Nozomi Nishiumi, Ayane Matsuo, Ryo Kawabe, Nicholas Payne, Charlie Huveneers, Yuuki Y. Watanabe, Yuuki Kawabata

## Abstract

Animal-borne accelerometers are effective tools for quantifying the kinematics of animal behaviors, such as swimming, running, and flying, under natural conditions. However, quantifying burst movements of small and agile aquatic animals (e.g., small teleost fish), such as during predatory behavior, or while fleeing, remains challenging. To capture the details of burst movements, accelerometers need to sample at a very high frequency, which will inevitably shorten the duration of the recording or increase the size of the device. To overcome this problem, we developed a high-frequency acceleration data-logger that can be triggered by a manually-defined acceleration threshold, thus allowing the selective measurement of animal burst movements. We conducted experiments under laboratory and field conditions to examine the performance of the logger. The laboratory experiment using red seabream (*Pagrus major*) showed that the new logger could measure the kinematics of their escape behaviors (i.e., body beat cycles and maximum acceleration values). The field experiment using free-swimming yellowtail kingfish (*Seriola lalandi*) showed that the loggers trigger correctly (i.e., of the 18 burst movements, 17 were recorded by the loggers). We suggest that this new logger can be applied to measure the burst movements of various small and agile animals, whose movements may be otherwise difficult to measure.

## Introduction

Animal-borne accelerometers have been used to estimate energy expenditure (Murchie et al., 2011; Payne et al., 2011), activity patterns (Kawabe et al., 2004; Payne et al., 2016; Sato et al., 2007), and specific behaviors such as the feeding, mating, and spawning (Føre et al., 2011; Tsuda et al., 2006; Watanabe and Takahashi, 2013; Whitney et al., 2010) of aquatic animals. In general, accelerometers record acceleration in a continuous manner at a defined frequency (e.g. 1–100 Hz) or record a defined time-average of the acceleration, either digitally stored or transmitted (Cooke et al., 2016). Subsequently, these acceleration data are often transformed to various components (e.g., dynamic and static accelerations) to estimate the energy budgets and activity, or to carry out classification into more detailed behaviors (Gleiss et al., 2011; Shepard and Wilson, 2008; Tanaka et al., 2001; Wilson et al., 2006).

Most previous studies have focused on routine behaviors such as cruising, gliding, and resting. Thus, the sampling frequencies of the accelerometers are typically below 32 Hz (Kawabe et al., 2004; Murchie et al., 2011; O'Toole et al., 2010; Tsuda et al., 2006). However, such low sampling frequencies cannot be used to measure the detailed burst movement dynamics of small and agile animals (e.g., teleost fish), which occur over short time scales (i.e., in the order of 100 ms), despite the fact that these burst movements include ecologically important behaviors such as escape responses and feeding strikes. The *in situ* measurements of such behaviors would provide novel insights into the movement performance, energy expenditure, and survival strategies of animals in complex natural habitats.

Recently, Broell et al. (2013) demonstrated that an accelerometer with a sampling frequency of at least over 30 Hz (ideally, 100 Hz) is required to identify the escape responses and feeding strikes of the sit-and-wait predator *Myoxocephalus polyacanthocephalus*. Moreover, accelerometers with a sampling frequency of 200 Hz have been demonstrated as useful in distinguishing the feeding behavior of trophic generalist fish on different prey types (Horie et al., 2017; Kawabata et al., 2014). These studies clearly show that the high-frequency accelerometers are useful in measuring the burst movements of agile animals; however, such high frequency sampling rapidly consumes electricity and memory, which inevitably shortens the duration of the recording, or increases its size (e.g., 10 hours in 100 Hz and 135 mAh, ORI400-D3GT from Little Leonardo Co., Tokyo, Japan). Thus, applying high-frequency accelerometers to field studies remains a challenging task.

To overcome this problem, we developed a data-logger, which selectively records acceleration signals based on a manually-defined threshold (Event logger), which reduces electricity and memory requirements, and thus enables us to measure the burst movements of animals for a relatively long period (e.g., 5-day battery life with a sampling frequency of 500 Hz and 10 burst movements per day) despite its small size (7.7 g in air). A similar selective recording system was used to measure the predatory behavior of the piscivorous pike *Esox lucius* in a laboratory setting (Van Deurs et al., 2017). However, the details of the logger system, the measurement performance of the logger, and its practicality for field measurements were not provided. In the present study, we describe the selective recording system of the Event logger, present results from the laboratory performance test conducted on the escape response of the red seabream *Pagrus major*, and show the results of the field performance test conducted on the yellowtail kingfish *Seriola lalandi*.

## Materials and Methods

### Event logger system

The Event logger consists of two different types of 3-axis accelerometers. One accelerometer detects the threshold excess (Detection accelerometer) and the other one records data (Recording accelerometer), as shown in Fig. S1a. The Detection accelerometer is continuously active, while the Recording accelerometer is inactive unless the Detection accelerometer detects any burst movements signals (i.e., exceeding a set threshold). The Detection accelerometer measures accelerations in the rate of 400 Hz. One absolute value is manually set as a threshold (any value from 1.00 to 3.95 *g*), and applied to the absolute values of all 3-axes accelerations. Once the Recording accelerometer becomes active, it records data for a manually set time period (any time period, in the order of 1 s). Then, it reverts to being inactive. If the acceleration exceeds the threshold when the Recording accelerometer is still active, the Detection accelerometer ignores it. The measurement range of the Recording accelerometer is ± 16 *g* with 16 bit resolution, and its sampling frequency can be set manually from 1 to 1,000 Hz. The battery life of the Event logger depends on the activation frequency and the recording period per activation (e.g., 5-day battery life with 10 activations per day, and a recording of 10 s per activation). The size, mass, and discharge capacity of the Event logger is 29 × 11 × 15 mm, 7.7 g, and 85 mAh, respectively (Fig. S1b). For interested users, the Event logger is available at http://www.biologging-solutions.com.

### Calibration of Detection accelerometer in the Event logger

Inherently, accelerometers have slightly different raw values among units, and the bias level can drift by the process of production, temperature fluctuation, large shock, etc. Therefore, the accelerometers need to be calibrated in advance. We calibrated the Recording accelerometer of the Event logger by referencing the gravitational acceleration and calibrated the Detection accelerometer by the method described below.

To find the formula for calibrating the Detection accelerometer, we examined the correspondence between actual acceleration values and the occurrence of the Event logger’s activation. The Event logger was attached to a similar-sized conventional 3-axis acceleration logger, which was continuously active (hereafter, Reference logger; 29 × 11 × 15 mm, 7.2 g, Biologging Solutions Inc., Tokyo, Japan). The sampling frequencies of both loggers were set to 500 Hz. These two loggers were packaged and thrusted, by hand, at various intensities (approximately 1–6 *g*). We set four different threshold values, namely 1.6, 2.4, 3.2, and 3.95 *g*, in the Event logger, and thrust 50 times in each threshold value. Logistic regression analysis was used to obtain the calibration formula. The occurrence of the Event logger’s activation was designated, in binary, as 1 (activated) and 0 (not activated); these values were used as the objective variable. The set threshold and maximum acceleration values, recorded by the reference logger, were considered as explanatory variables. In addition, we calculated the time lag between the time when the acceleration exceeded the threshold, and the time when the recording was initiated.

### Experiment 1: Laboratory performance test using red seabream

To examine the measurement performance of the logger, we attached the logger on *P*. *major* in a tank and measured its escape response.

Four *P*. *major* were obtained from a local fish hatchery and transported to the Institute for East China Sea Research at Nagasaki University, Japan. The fish were held in 500 L circular polyethylene tanks (100 cm diameter × 75 cm height), with an aeration apparatus and flow-through seawater at a temperature of 19.9–24.2 °C. The mean body mass and total length of the fish were 2.50 kg (range: 1.96−3.05 kg) and 52.5 cm (range: 49.9−54.8 cm), respectively.

We attached the logger package incorporating the Event logger and the reference logger onto the *P*. *major*. The fish were first anaesthetized using 0.1% 2-phenoxyethanol; then, the logger was attached using two wiry plastic strings and two 2-cm round stainless washers, which served as anchors. The strings were first inserted through the tag, and then through the anterior dorsal musculature, after passing through two syringes. Finally, they were anchored in place by round stainless washers. The tagging procedure never exceeded 1 min. The loggers’ x, y, and z axes were aligned to the lateral (rightward), longitudinal (forward), and vertical (upward) coordinates of the fish body, respectively. The sampling frequencies of both loggers were set to 500 Hz. The threshold value of the Event logger was set to 2.0 *g*, since the excess of this value rarely occurred during the routine movements of other fish species, namely, *Epinephelus ongus* (Kawabata et al., 2014) and *Seriola quinqueradiata* (Noda et al., 2013). The recording period of the Event logger was set to 5 s per activation.

The experiment was performed in a 3,000 l circular FRP tank (193 cm diameter × 73 cm height), which was filled with seawater to a depth of 30 cm. The fish were individually introduced into the tank and were allowed to acclimate for approximately 20 h. Then, we plunged a hand net into the water, near the fish, and induced the escape response. For each fish, the escape response was elicited 8–10 times, in 30-minute intervals, and a total of 34 trials were carried out across the four fish. To determine the timing at the initiation of the escape response, the fish movements were simultaneously recorded dorsally, by a high-speed video camera (HAS-L1; Detect Co., Tokyo, Japan), at 500 frames s^−1^. The water temperature was 20.6–22.5 °C.

#### Data analysis

We used 11 of the 34 trials, while the remaining 23 trials were omitted either because the fish did not show any escape response against the hand net, the response was disturbed by the tank wall or intensive waves, or the hand net obscured the fish body.

Maximum and minimum acceleration values, and oscillation cycles, are important variables for estimating locomotor performance and categorizing animal behaviors (Broell et al., 2013; Kawabe et al., 2004). Since latency existed between the initiation of escape response and the recording initiation of the Event logger, we examined whether the latency was short enough to precisely measure these variables. To measure the exact timing of escape response initiation and the recording initiation of the Event logger, we synchronized the acceleration signals of the Event logger to those of the reference logger by finding the optimal time difference with the least squares method. The escape response of fish consists of three distinct stages based on body bends (Weihs, 1973). Peak acceleration usually occurs during the initial bend, or stage 1 of the escape response (Domenici, 2009). Therefore, the peak acceleration timings during stage 1 were used in the analysis.

To specify the sampling frequency required for the precise measurement of the above variables, we examined the effect of sampling frequency on the measured variables. The accelerations of different sampling frequencies (1–500 Hz) were obtained by downsampling the 500 Hz acceleration signals from the Event logger. The maximum and minimum values of the downsampled accelerations were then compared to those of the raw 500 Hz accelerations. We also calculated the minimum sampling frequency required for detecting the oscillation cycles of the acceleration signals, which usually reflects the tail beat frequencies of fish (Kawabe et al., 2003). At least two points within an oscillation wavelength are necessary to measure the oscillation cycle; thus, we regarded the required sampling frequency as reciprocal to one half of the wavelength. In this analysis, we used x-axis accelerations, since tail beats produce mainly lateral accelerations (Kawabe et al., 2003),

### Experiment 2: Field performance test using yellowtail kingfish

We examined the accuracy of the Event logger’s recording initiation system under natural conditions by attaching an Event logger and reference logger to free-ranging *S. lalandi* and comparing the obtained acceleration values. This experiment was conducted off the Neptune Islands Group (Ron and Valerie Taylor) Marine Park, Australia (S35° 14′, E136° 04′) from October to November 2015.

Three fish (98, 99, and 102 cm TL) were caught by hand line and tagged on board. Fish were placed on a rubber mat and their head was covered with a wet towel while the gills were continuously ventilated with a saltwater hose. In compliance with local animal ethics procedures, fish were not anaesthetized. We attached the logger package consisting of a radio tag, an Event logger, a reference accelerometer logger (ORI400-D3GT, 12 mm diameter × 45 mm length, 9 g, 135 mAh; Little Leonardo Co., Tokyo, Japan), and a time-scheduled release mechanism (Watanabe et al., 2004) to each fish. The tagging procedure of the logger package was similar to the Experiment 1. The fish were released promptly after the tagging, and the tagging procedure never exceeded 3 min. The sampling frequency, threshold value, and per-activation recording period of the Event logger, were set to 500 Hz, 2.0 *g*, and 10 s, respectively. The reference logger was set to continuously measure 3-axis accelerations at 20 Hz. Approximately 45, 37, and 18 hours after release, the logger packages popped off from the fish, and emerged afloat for recovery.

Two of the three Event loggers worked properly and were thus used in the data analysis. We synchronized the acceleration signals from the Event logger to those from the reference logger using the least squares method. Subsequently, we examined whether the Event loggers were accurately activated, by comparing the activation events with the threshold excess in the acceleration signals obtained from the reference loggers. The maximum and minimum acceleration values, and minimum wavelength obtained through the Event loggers were compared to those obtained through the reference loggers.

## Results

### Calibration of Detection accelerometer in the Event logger

In all four set threshold values, the maximum accelerations were higher in the trials where the Event logger was activated compared to trials where the Event logger was not activated. The acceleration range at the 1.6 and 2.4 *g* thresholds, however, slightly overlapped between the activated and non-activated trials (Table S1). The calibration formula was obtained by a logistic regression and successfully classified 95.5 % (191/200) of thrust events (Fig. S2). The mean time lag between the instance when the acceleration exceeded the threshold and the instance when the recording was initiated was approximately 1.11 × 10^−2^ s (Table S1).

### Experiment 1: Laboratory performance test using red seabream

The Event logger initiated recording at 1.64 × 10^−2^ ± 1.21 × 10^−2^ s (mean ± SD) after the initiation of the escape response. In all of the 11 trials, the initiation occurred before the first half of the stage 1 escape response, which indicated that the recording latency was short enough to measure the oscillation cycle of the acceleration signals with precision. Successful recording rates of maximum and minimum acceleration values by the Event logger in the stage 1 escape response are shown in Fig. 1. Of the 11 trials, 11 maximum (100 %) and six minimum (55 %) x-axis accelerations, 10 maximum (91 %) and 5 minimum (45 %) y-axis accelerations, and nine maximum (82 %) and two minimum (18 %) z-axis accelerations were successfully recorded by the Event logger. Depending on which of the two came first, the maximum or the minimum acceleration values occurred at 8.55 × 10^−3^ ± 2.54 × 10^−3^ s (mean ± SD) after the escape response initiation.

**Fig. 1.**
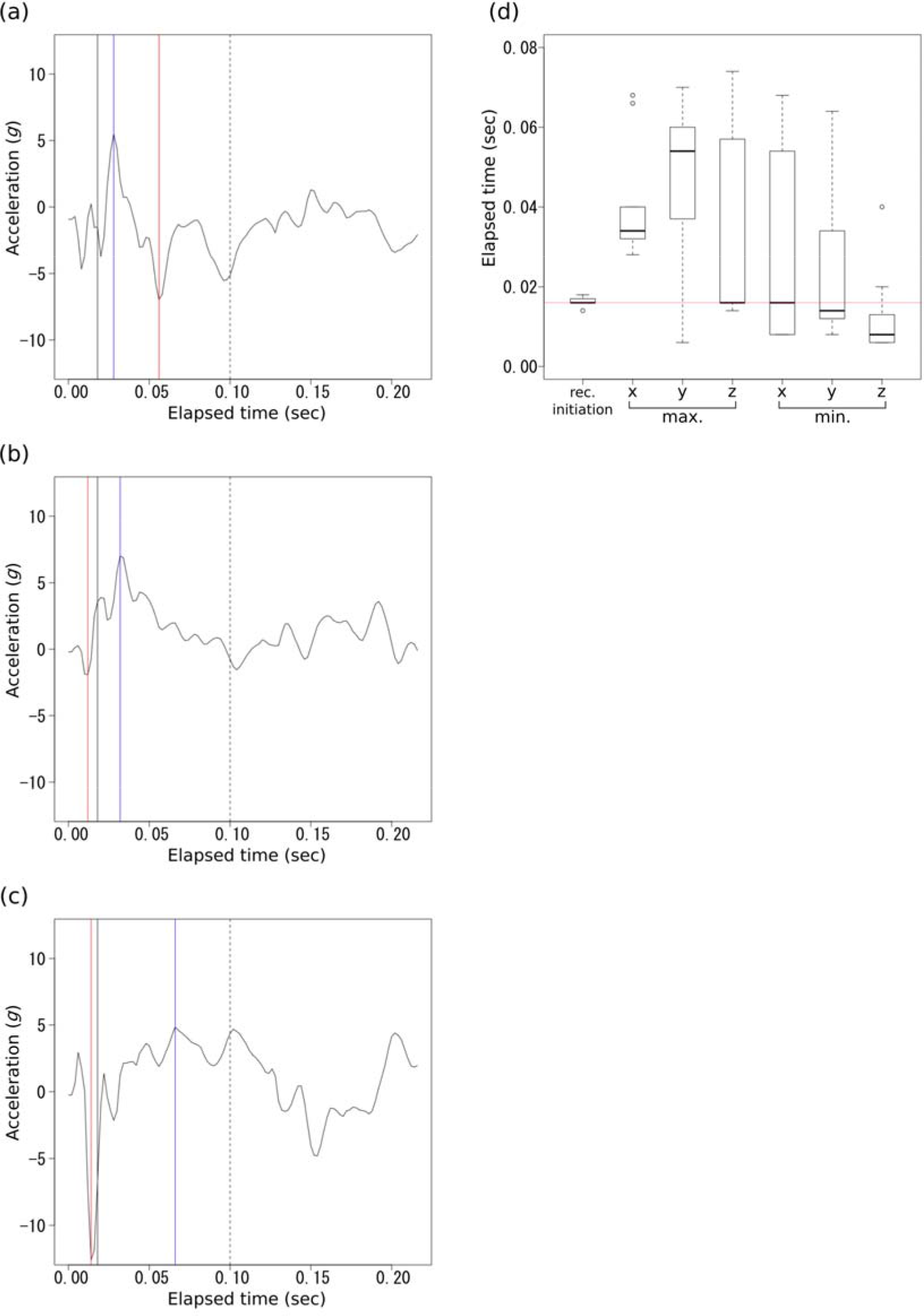
(a-c) Typical acceleration signals (a, x-axis; b, y-axis; c, z-axis) during the escape response of red seabream. The black solid and dashed lines indicate the initiation of recording and the end of the stage 1 escape response, respectively. Blue and red lines indicate the instances when acceleration reached their maximum and minimum acceleration values, respectively. (d) Boxplot of record initiation latency, and the instances when acceleration reached maximum and minimum acceleration values. The boxes, upper and lower whiskers indicates IQR, maximum value or Q_3_ + 1.5 IQR, and minimum value or Q_1_ − 1.5 IQR, respectively. The red line indicates the median latency of record initiation

The absolute values of maximum and minimum accelerations became smaller as the sampling frequency decreased. The median value of the downsampled acceleration signals became less than 95% of the original values at 50.0–166.7 Hz and less than 75 % of the original values at 22.7–100.0 Hz (Figure 4). The minimum wavelength median of the acceleration in the x axis was 4.8 × 10^−2^ s, and the required sampling frequency for detecting the oscillation cycle was estimated at 41.7 Hz.

### Experiment 2: Field performance test using yellowtail kingfish

Each of the acceleration signals recorded by the two reference loggers, namely, packages 1 and 2, exceeded the set threshold nine times each. The Event logger of package 1 was activated in all nine excesses, while that of package 2 was activated in eight of the nine excesses. The maximum acceleration value of the remaining one excess was 2.42 *g*. In addition to the nine and eight activations, both Event loggers were activated another five times each, when the acceleration measured by the reference loggers surged, but did not exceed the threshold (mean 1.72 ± 0.20 *g*, N=10, Fig. 3). With the exception of the maximum x-axis acceleration values and minimum x- and y-axes values in package 2, the maximum and minimum values obtained through the Event loggers were larger and smaller, respectively, than those obtained via the reference loggers (Table S2).

## Discussion

The experiment with red seabream showed that the Event logger recorded accelerations during most parts of the escape responses, with sufficient temporal resolutions.
Although there was latency in initiating the recording, which prevented accurate recordings of minimum acceleration values during the stage 1 escape response, the logger successfully recorded the maximum acceleration values and oscillation cycles with high accuracy. The difference in successful recording rates, between maximum and minimum acceleration values, was related to the asymmetrical waveform of the acceleration signals (see Fig. 1). The asymmetrical waveform may be due to the gravitational, propulsive and centripetal accelerations, and to the logger attachment site (i.e., left side of the fish body); however, further research is required to explicitly clarify the cause of this asymmetry.

The latency in the record initiation was a combination of time lag from escape initiation to threshold detection and time lag from threshold detection to record initiation. Although the former time lag can be modified by a set threshold value, the latter time lag is mechanically fixed as the mean of 1.11 × 10^−2^ s (Table S1). Because the first peak value (maximum or minimum) occurred at 8.55 × 10^−3^ s after the escape response initiation, recording both maximum and minimum values during the stage 1 escape response of this species was difficult with the present system. Nonetheless, the time to peak acceleration should be species- and behavior-specific; thus, the latency could be short enough to record both maximum and minimum acceleration values for other studies. Additionally, the latency will be shorter as sensor technology continues to advance.

The escape response of red seabream included abrupt acceleration changes and thus more than 166.7 Hz of sampling frequency was required to measure the exact peak accelerations, and more than 41.7 Hz for estimating tail beats. This result is consistent with the conclusion by Broell et al. (2013), by which accelerometers with a sampling frequency of more than 30 Hz are required to identify the escape and predatory behaviors of fish.

The results of the field study show that the Event logger was accurately activated in response to the threshold excess. The Event logger failed to detect one threshold excess; however, this could have been caused by a slight positional difference between the Event logger and the reference logger. The Event logger was also activated several times, when the acceleration measured by the reference logger did not exceed 2.0 *g*. Although these activations could be caused by false detections by the Detection accelerometer (see Table S1), it is more likely that the activations are related to the 400 Hz samplings of the Detection accelerometer, which allows the detection of momentary high acceleration with higher probabilities than the 20 Hz samplings of the reference logger (See Fig. 2).

**Fig. 2.**
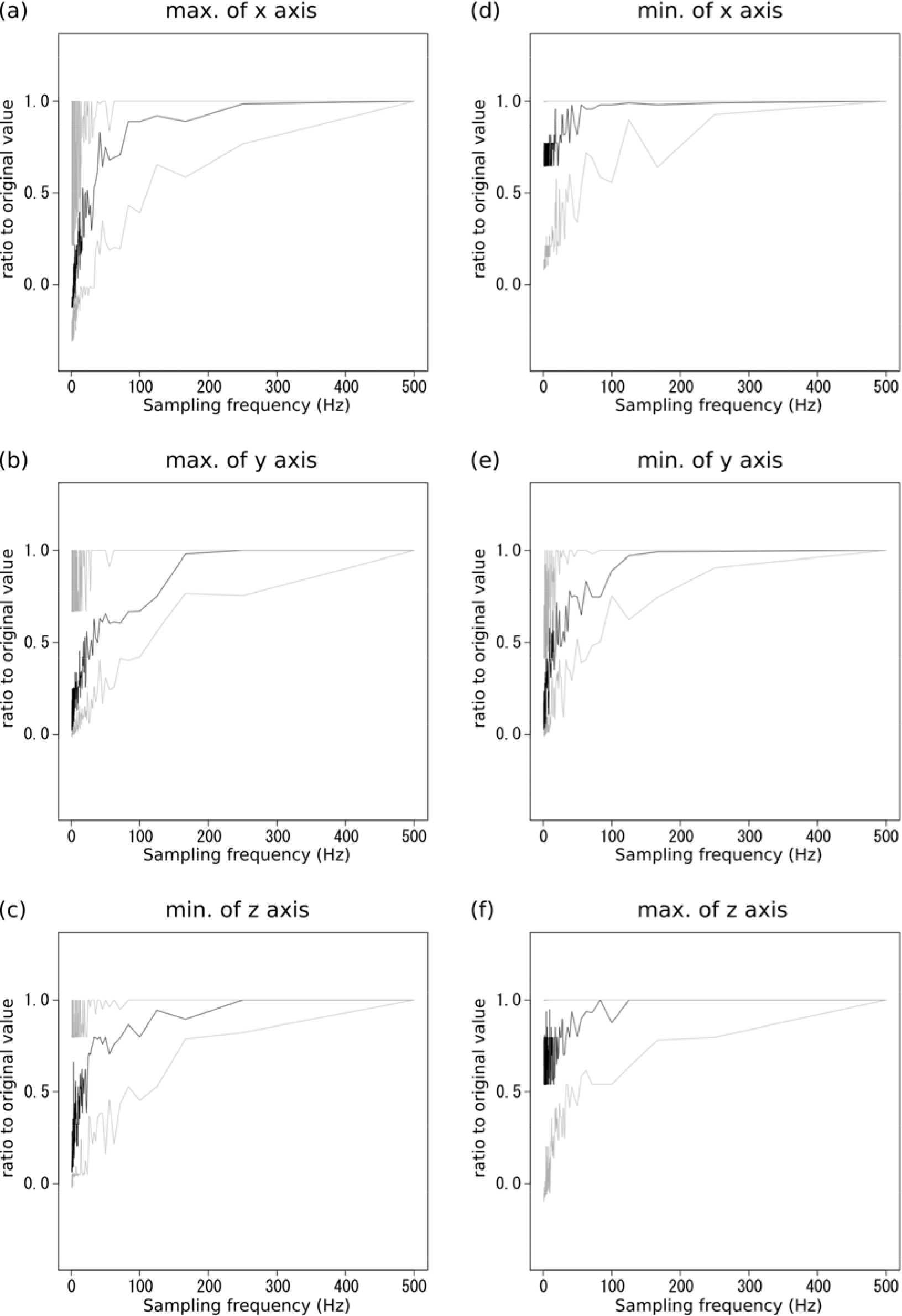
The effect of sampling frequency on measured maximum and minimum acceleration values during the escape response. The vertical axes show the ratio of the downsampled acceleration data to the original 500 Hz acceleration data. The solid black line and upper and lower gray lines indicate median, maximum and minimum values, respectively

**Fig. 3.**
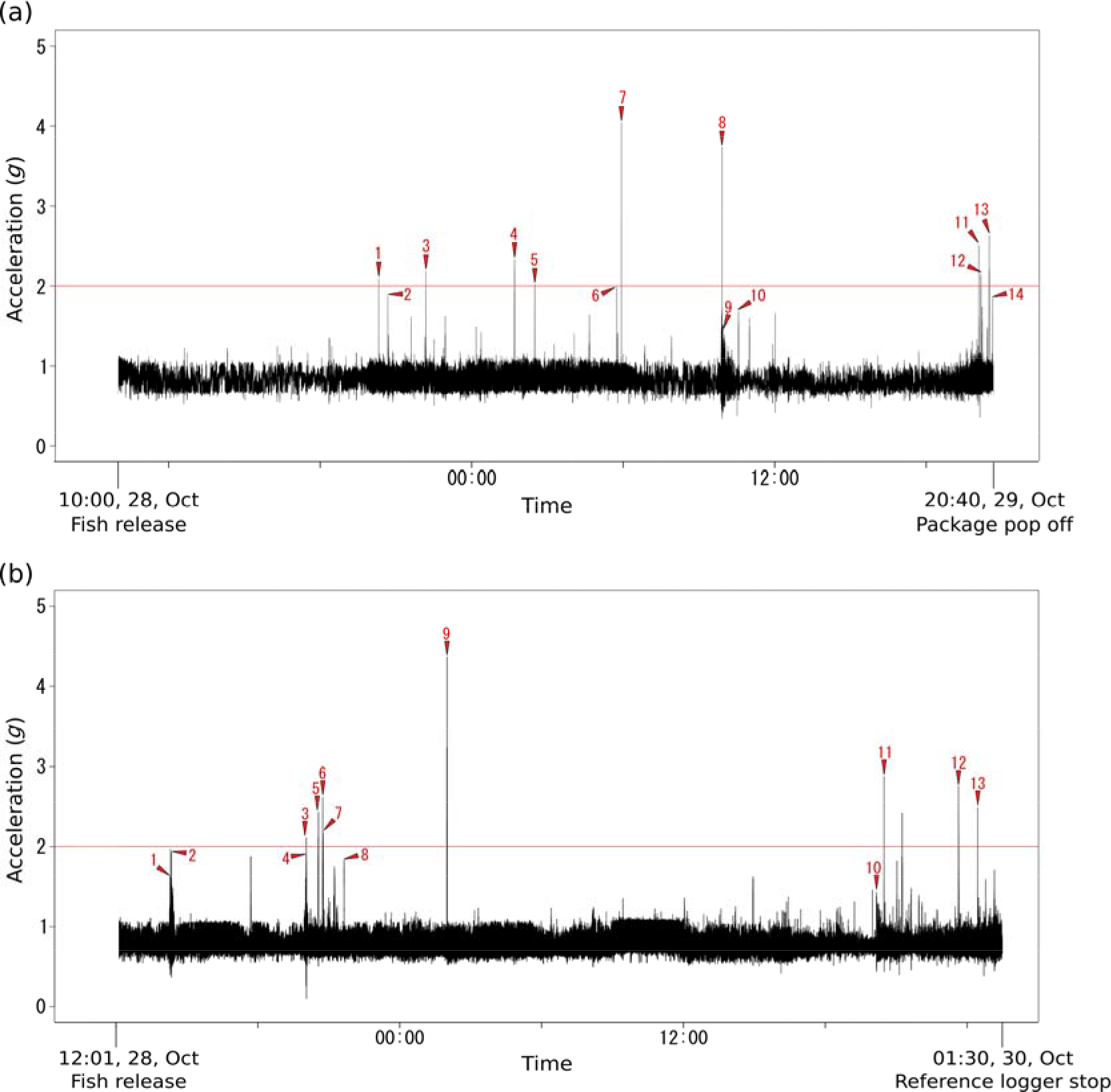
Activations of the Event loggers attached onto the two free-ranging yellowtail kingfish (a, b). The activation events (red arrows) are shown against the maximum absolute values of the 3-axis accelerations recorded by the 20 Hz reference loggers

The Event logger system is similar to on-board data processing, in the sense of discarding unnecessary data before recording. Generally, on-board data processing operates by summarizing the information inside the tags, which allows us to save battery and minimize memory usage. Previous studies demonstrated that the acceleration data could be processed into activity level (Payne et al., 2016) or occurrence of specific behaviors, such as opening jaws, stroking flippers, and borrowings (Adachi et al., 2014; De Almeida et al., 2013; Naito et al., 2013). However, few on-board processing methods have been reported to determine the types of burst movements, possibly due to the low sampling frequencies of accelerometers, and due to the limited numbers of established processing algorithms (but see (De Almeida et al., 2013; Horie et al., 2017). In addition, such processing would limit the scope of analysis, since it cannot provide the various kinematic variables simultaneously. In contrast, the Event logger works with a simple algorithm, which records certain raw acceleration data with a high sampling frequency, and thus allows post-hoc analysis from various kinematic aspects. Such raw high-frequent acceleration data would be especially useful for exploring burst behaviors, which have not been studied well. Therefore, we believe that this new logger will contribute to the exploration of burst movements in various animals, and to the further development of animal-borne accelerometer methods.

## Acknowledgments

We are grateful to G. N. Nishihara, I. Nakamura, and H. Kimura for their assistance in experiment 1, and to W. Robbins, L. Meyer, S. Whitmarsh, A. Schilds, L. Nazimi, S. Payne, R. Hall, M. Ward, and R. Mulloy for assisting with experiment 2.

## Author contributions

Y.K. developed the data logger and conceived the study. A.M. and Y.K. conducted the calibration experiment. N.N., A.M., and R.K. conducted the laboratory experiment with the red seabream. N.P., C.H., and Y.Y.W. conducted the field experiment on the yellowtail kingfish. N.N. and Y.K. analyzed the data. N.N. and Y.K. drafted the manuscript with critical input from the other authors.

## Funding

This study was funded by Grants-in-Aid for Scientific Research, JSPS, Japan to Y.K. (25870529), R.K. (26450263 and 16H05795), and Y.Y.W. (25850138), Sumitomo Foundation to Y.K. (153128), and Sustainable Aquatic Food and Environment Project in the East China Sea, MEXT, Japan to Nagasaki University. N.P. was funded by a JSPS Postdoctoral Fellowship for Research in Japan. The fieldwork for the yellowtail kingfish component was funded by the Neiser Foundation, Fox Shark Research Foundation, and Nature Films Production.

## Ethical approval

Animal care and experimental procedures were approved by the Animal Care and Use Committee of the Institute for East China Sea Research, Nagasaki University (Permit no. ECSER15-13), in accordance with the Regulations of the Animal Care and Use Committee of the Nagasaki University, and Animal Care and Ethics Committee of the University of New South Wales (Permit no. 15/126B). The field work was carried out under the Department of the Environment, Water, and Natural Resources (DEWNR) scientific research (Permit no. M26292), and Marine Parks (Permit no. MR00047).

